# Phenotype switching in a global method for agent-based models of biological tissue

**DOI:** 10.1101/2022.08.22.504898

**Authors:** Daniel Bergman, Trachette L. Jackson

**Affiliations:** Department of Mathematics, University of Michigan, Ann Arbor, MI 48109, USA

## Abstract

Agent-based models (ABMs) are an increasingly important tool for understanding the complexities presented by phenotypic and spatial heterogeneity in biological tissue. The resolution a modeler can achieve in these regards is unrivaled by other approaches. However, this comes at a steep computational cost limiting either the scale of such models or the ability to explore, parameterize, analyze, and apply them. When the models involve molecular-level dynamics, especially cell-specific dynamics, the limitations are compounded. We have developed a global method for solving these computationally expensive dynamics significantly decreases the computational time without altering the behavior of the system. Here, we extend this method to the case where cells can switch phenotypes in response to signals in the microenvironment. We find that the global method in this context preserves the temporal population dynamics and the spatial arrangements of the cells while requiring markedly less simulation time. We thus add a tool for efficiently simulating ABMs that captures key facets of the molecular and cellular dynamics in heterogeneous tissue.

**Author summary:** Agent-based models (ABMs) are an important tool for understanding how cells and molecular compounds interact to produce complex, emergent behavior. The principal feature of ABMs that set them apart from other types of models is their ability to capture the diversity of cells in a tissue. However, this feature comes at the cost of long simulation times, reducing the ability to apply the findings of these models to improve our understanding of living organisms. We present here a means of simulating ABMs using a more efficient method, called the global method, when the cells are undergoing discrete, phenotypic changes in response to molecular cues. We demonstrate that the global method preserves the key features of the ABM while performing simulations much faster. This allows for more efficient testing of biological hypotheses in a mathematical framework that captures key facets of the diversity in biological tissue.

## Introduction

Agent-based models (ABMs) are a fixture in mathematical biology, having grown in use and significance dramatically over the past two decades. Their ability to capture biological processes across spatial and temporal scales make them well-suited for exploring the complex dynamics of living tissue [1]. A critical feature of this interplay is the incredible level of phenotypic diversity of the cells that constitute biological tissue. This heterogeneity has many sources: genes, cellular specialization, environmental factors, etc. ABMs must be constructed to contain sufficient levels of heterogeneity so that the insights gleaned from them can be reliably applied to the biological system they attempt to describe.

The trade-off is that ABMs are computationally expensive when compared to other modeling approaches such as differential equation models. This typically limits ABMs run on a desktop workstation to around 10^6^ cells [2], despite one cubic centimeter of tissue containing upwards of 10^8^ cells [3]. To push ABMs to the giga-scale [4], it is necessary to develop techniques that can reduce the cost of simulating ABMs.

Many modelers have been discussing and putting forth strategies for speeding up ABM simulation time or tasks requiring simulations of ABMs. These techniques include modeling a small-but-relevant region of the tissue [5], partitioning the microenvironment into compartments without internal spatial resolution [6], and others. Targeting the specific task of parameters estimation, [7] laid out an efficient means for performing Bayesian inference on an ABM. In [8], we developed and analyzed what we refer to as the global method for handling molecular dynamics in an ABM, and this method resulted in orders of magnitude speedup. This method is best suited for ABMs that require simulating reactions at the cell surface and/or intracellular signaling pathways. The traditional way of simulating such ABMs, which we call the local method, is to solve a system of ordinary differential equations for every cell at every time step. The global method reduces the computational time compared to the local method by averaging the molecular state variables across cells in a region of the microenvironment before solving the governing differential equations and applying the result to those cells. In [8], we explored two model systems in which these molecular dynamics resulted in continuous changes to cellular dynamics by modulating proliferation, apoptosis, and symmetric division rates. Among the many questions this left open regarding the applicability of the global method, a critical one was how it performed when the molecular level caused discrete changes at the cellular level. That is, can the global method perform well when the molecular level causes phenotypic changes in cells such as a transition between states?

One of the most fundamental features of life is the need for oxygen. In the absence of oxygen, cells can undergo various changes to adapt to the stress induced by a hypoxic environment [9–12]. In other words, cells often change states based on the presence or absence of oxygen. Because of the ubiquity of this process in biological tissue, we choose it to explore how the global method performs in the context of phenotype switching.

We will look at cells growing near vasculature that generates an oxygen gradient as oxygen extravasates, diffuses and degrades, exchanges with the cells, and is finally metabolized by the cells. Cells that acquire sufficient oxygen, will remain in a proliferative state whereas those that do not will transition to a quiescent state that is non-proliferative. We will also explore how the global method works when the quiescent cells can act with a level of agency by moving towards the oxygen source in an effort to return to the proliferative state. Finally, we also allow the proliferating cells to cluster together by following a gradient of a second substrate, called a quorum factor, that is secreted by all cells.

We also introduce a hybrid method that combines pieces of each method by using the local method to solve cell-independent molecular dynamics and then using the global method to solve cell-dependent dynamics. The hybrid method thus focuses on the greatest computational savings provided by the global method while retaining more molecular spatial heterogeneity than the global method.

## Methods

### Agent-based model

We use an on-lattice ABM with reflecting boundary conditions to explore how the global method performs when phenotypic switching can occur throughout a simulation. There are two types of agents in this model: proliferating and quiescent cells. Proliferating cells proliferate, move, and die at fixed rates given in Table 1. Quiescent cells do not proliferate and their death and movement rates are orders of magnitude slower than those for proliferating cells (Table 1).

**Table 1.** Parameter values.

| Name | Description | Value |
| --- | --- | --- |
| $p$ | proliferation rate of proliferating cells | $2 \text{ d}^{-1}$ |
| $d$ | apoptosis rate of proliferating cells | $2 \times 10^{-3} \text{ d}^{-1}$ |
| $m$ | movement rate of proliferating cells | $2 \mu\text{m min}^{-1}$ |
| $d_Q$ | death rate of quiescent cells | $2 \times 10^{-5} \text{ d}^{-1}$ |
| $m_Q$ | movement rate of quiescent cells | $0.2 \mu\text{m min}^{-1}$ |
| $C_{\text{sys}}$ | concentration of oxygen in systemic circulation | 38 mmHg |
| $f$ | fluid exchange rate of oxygen across capillary walls | $20 \text{ min}^{-1}$ |
| $D$ | diffusion coefficient of oxygen | $1 \times 10^5 \mu\text{m}^2 \text{ min}^{-1}$ |
| $\lambda$ | degradation rate of oxygen | $0.1 \text{ min}^{-1}$ |
| $h$ | spatial discretization on lattice | 20 $\mu\text{m}$ |
| $u$ | uptake rate of oxygen | $10 \text{ min}^{-1}$ |
| $s$ | secretion rate of oxygen | $10 \text{ min}^{-1}$ |
| $k_{\text{aer}}$ | rate of aerobic respiration | $1 \times 10^{-6} \text{ mmHg}^{-5} \text{ min}^{-1}$ |
| $I_{\text{thresh}}$ | quiescence threshold | 5 mmHg |
| $I_{\text{min}}$ | threshold below which the quiescence rate is at a constant, maximal rate | 2.5 mmHg |
| $I_{\text{max}}$ | threshold above which the quiescence rate is 0 | 7.5 mmHg |
| $q_{\text{max}}$ | maximal quiescence rate | $0.01 \text{ min}^{-1}$ |

The simulation advances by a Gillespie algorithm, randomly deciding on the next time step given the sum of all rates in the model. The one event for that time step is chosen randomly with weights given to each event for each agent based on the rate of that event. The model advances forward in time through such time steps until the next randomly chosen time step moves the simulation past the predetermined end time, in which case no cell events are performed.

### Molecular dynamics

Since the molecular dynamics are assumed to be much faster than the cellular dynamics, the model uses a quasi-equilibrium assumption during the above event selection process. That is, the molecular dynamics are updated at regular intervals and are assumed constant throughout one such interval. The length of these intervals is approximately Δ*t*_mol_ as explained below.

The main substrate in this model is oxygen and there are four differential equations that it obeys: pharmacokinetics (PK), diffusion, cellular exchange, and intracellular signaling, i.e. cellular metabolism. We also include a second substrate, quorum factor, in some simulations that only undergoes diffusion and cellular exchange. These are solved in this order, repeating as necessary until the next time to update the cellular dynamics. One iteration through this repetition updates each individual reaction by a set time step. Within that update, each reaction follows its own time step to advance with this time step chosen to achieve numerical stability. By fixing this reaction-specific time step, we can perform pre-computations that reduce the computational costs, paritcularly for the analytic solutions relying on a matrix exponential. The techniques used to solve each of these differential equations in the two methods is summarized in Table 2.

**Table 2.**
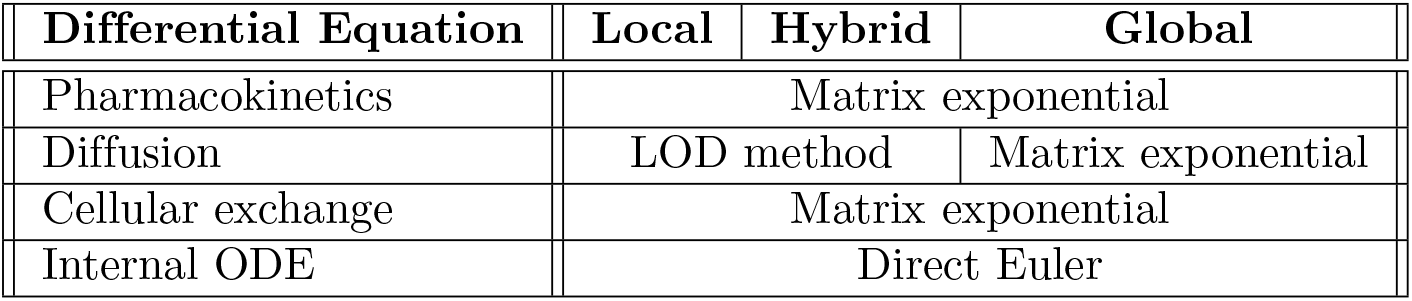
Solvers used for each differential equation in the methods.

We also make use of a hybrid method. This method maintains the full spatial resolution of the substrate in the microenvironment for the PK dynamics and diffusion, using the same techniques as the local method to solve these as described below. It then applies the techniques of the global method to solve the cellular exchange and intracellular signaling differential equations.

### Regions: coarse-graining the microenvironment

Both the global and hybrid methods require partitioning the microenvironment into regions. That is, each lattice site is assigned to a single region (see Fig 1A). The average of the molecular state variables within each region will be used to solve the differential equations, rather than the specific concentrations at each lattice site and cell. The choice of regions can be made in any way, but different choices will lead to different levels of agreement with the local method. In this work, we shall make use of two choices of regions depending on the location of vasculature in the simulation. For the majority of our work, we shall assume that the vasculature is located at the bottom of the microenvironment, i.e., at *y*_min_ in 2D simulations and at *z*_min_ in 3D simulations. Under this vasculature assumption, regions will be layers stacked on top of one another, each one cell thick. We shall also consider the case in which vasculature surrounds the growing cells. In this case, regions will be chosen as concentric annuli around the center of the microenvironment in two dimensions. For three dimensions, we will use concentric spherical shells.

**Fig 1.**
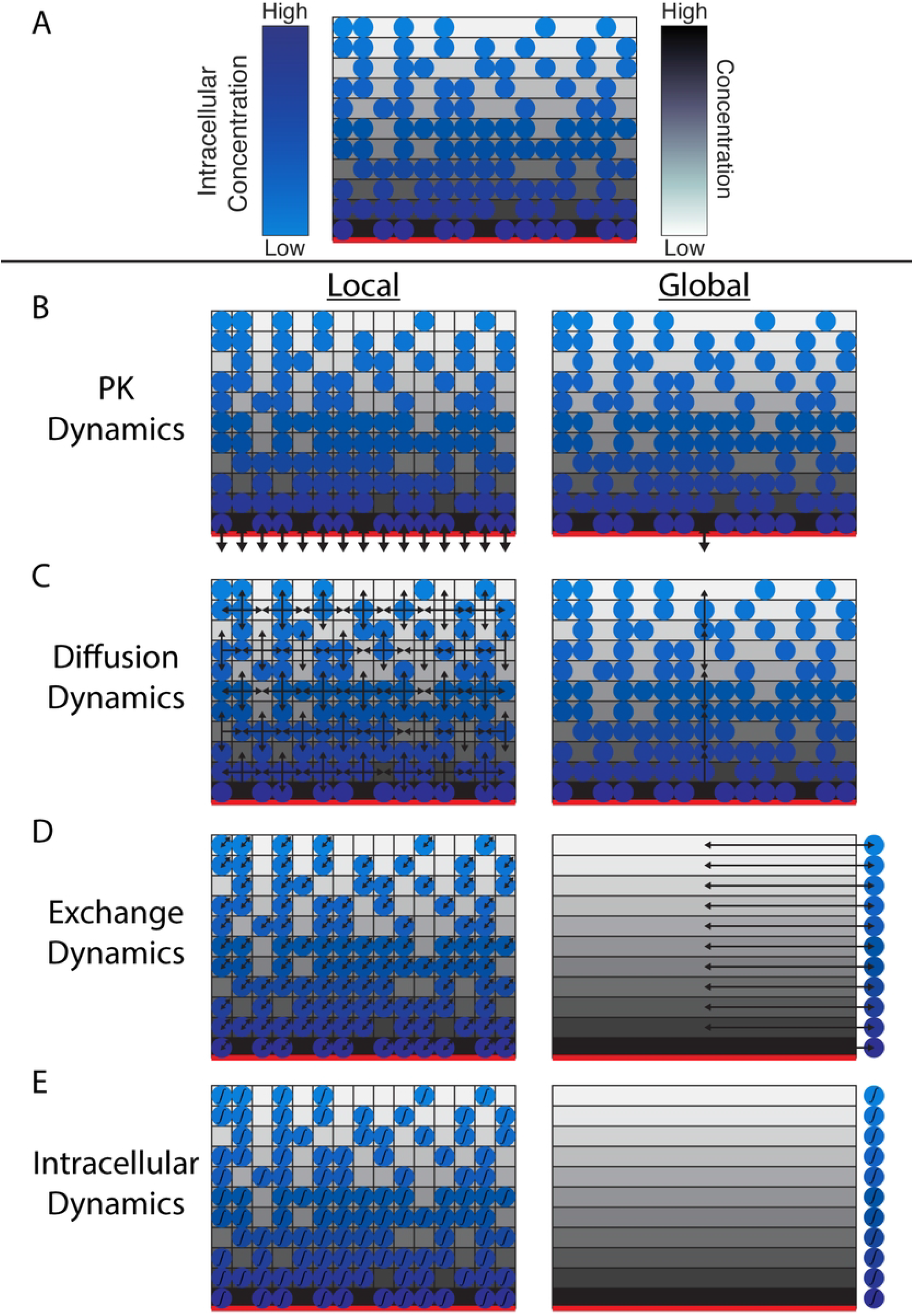
Schematic comparison of the two methods. A. Schematic of regions in the global method when the vasculature is at the bottom boundary. The red line along the bottom indicates the location of the blood vessel. The monochrome rectangles represent regions used in the global method. The discs represent agents in the model. The shading of each region indicates the average oxygen concentration within. The shading of each agent represents the internalized concentration. The bottom-most region is the only region containing perivascular lattice sites. All of the lattices sites in the bottom region are perivascular. B-E. Schematic comparison of how the local (left column) and global (right column) methods treat the four molecular-level dynamics. B-C. In the local/global methods, PK dynamics update perivascular sites/regions (B) and diffusion occurs between neighboring sites/regions (C). D-E. The local method solves the exchange and then intracellular dynamics for each cell individually. The global method solves these once per region.

In both assumptions about vasculature, the regions can be understood as subsets of lattice sites with equal access to oxygen. This is borne about when looking at simulations of the local method (Fig S1 Fig). When we introduce a quorum factor, first solve the ABM using the local method for the quorum factor dynamics because it is not clear *a priori* what regions could be constructed in which that substrate is approximately constant. After observing these dynamics, we then choose an appropriate region scheme to solve the quorum factor dynamics using the global method.

### Pharmacokinetics

Pharmacokinetics (PK) describes how the substrate can enter the microenvironment from outside the microenvironment (Fig 1B). Here, we assume that the body provides a constant source of blood and oxygen to the capillaries running through the microenvironment. That is, the circulation concentration of oxygen, *C*_sys_ is treated as a constant. We also assume that passive diffusion is responsible for oxygen crossing the capillary walls in either direction. Thus, the PK for this model is given by the single ODE in Eq 1 where *C* represents the concentration at any perivascular lattice site and *f* is a parameter controlling the rate of exchange of oxygen between the blood and interstitium.

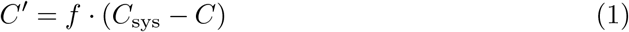

When using the local method, we solve this equation for each perivascular lattice site, which can be solved efficiently using the analytic solution and vectorization in MATLAB. In the global method, Eq 1 still holds, but only for those regions that are perivascular. We define regions so that either all of the lattice sites or none of the lattice sites in a region are perivascular (Fig 1A).

In most of our vasculature schemes, a lattice site being perivascular is a Boolean property of the given site, i.e., it is or is not perivascular. In our final example using a spherical shell of vasculature, this binary approach results in a non-uniform density of vasculature on the shell due to the incompatibility of a sphere and a lattice. In this example, we instead assign to each lattice site a non-negative value quantifying the volume of vasculature at that lattice site where a value of 0 signifies the lattice site is not perivascular and larger positive values indicate closer proximity to vasculature. Calling this value *p* for a given lattice site, we replace *f* in Eq 1 with *p* · *f*. For the global method, we average across a given region all the values of *p* assigned to each lattice point in the region. This average, 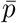, is used to replace *f* with 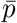 · *f* for this region.

### Diffusion

Diffusion describes the dynamics of the substrate within the microenvironment in the absence of cells (Fig 1C). We assume that once oxygen is in the microenvironment, it diffuses freely throughout and also undergoes spontaneous degradation. This degradation could be interpreted as metabolism by cell types other than those considered explicitly in the ABM. Thus, the diffusion equation is given by Eq 2

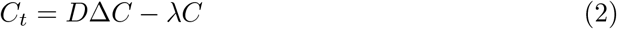

together with a no-flux, or Neumann, boundary condition. When using the local method, we solve this PDE with a locally one-dimensional (LOD) method. In the global method, however, we discretize the Laplacian using a standard stencil and use that to rewrite Eq 2 as a system of ODEs describing the average concentration of oxygen within each region. The resulting system for the oxygen concentrations in region *i, C*_*i*_, is given by Eq 3.

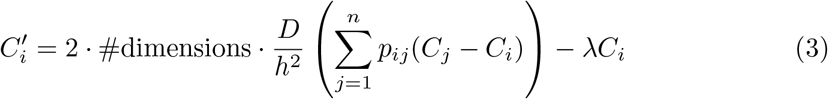

where #dimensions is the number of dimensions simulated in the ABM (we use both 2- and 3-dimensional simulations here), *h* is the spatial discretization of the lattice, and *p*_*ij*_ is the proportion of region *i* neighbors located in region *j*. In other words, weight the diffusion based on how much region *i* neighbors region *j*. The matrix *p*_*ij*_ need not be symmetric in general, but it is often sparse as most regions only neighbor a small subset of all regions. Eq 3 is linear and so the analytic solution can be readily used. Furthermore, since the entries in the defining matrix of this system of ODEs is fixed once the regions are decided, its matrix exponential for a given time step can be pre-computed and reused throughout the simulation.

### Cellular exchange

Cellular exchange describes how cells affect the concentration of the free substrate in the microenvironment as well as any uptake of the substrate into the cells (Fig 1D). Here, we assume that oxygen diffuses passively through cell membranes and that the oxygen inside the cell is well-mixed. Thus, we use Eq 4 to describe how the concentration in and around each cell changes due to cellular exchange. In this equation, *C* represents the concentration of oxygen in the microenvironment at the lattice site and *I* represents the internalized concentration of oxygen in the cell at that lattice site.

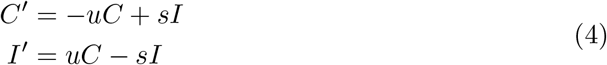

When using the local method, we solve this equation for each cell. Due to the linearity of this equation, we can even solve this equation using a matrix exponential. By using a fixed time step for solving this equation, then we can store this matrix exponential to be reused for the entire simulation.

In the global method, Eq 4 is used for each (region, type) pairing with the state variables representing the average concentrations within a region *i* occupied by cell type *k*. The same matrix exponential as in the local method can be used for each (region, type) pair. This then produces a solution to the extracellular concentration within region *i* for each cell type *k*, which must be combined to get a new average concentration in the region. To do this, we set *q*_*ik*_ to be the proportion of region *i* occupied by cell type *k* and let *C*_*ik*_(*t*) be the extracellular component of the solution on *t* ∈ [0, *t*_*f*_]. Note that the *q*_*ik*_ will vary throughout the simulation due to cellular events such as proliferation, apoptosis, and movement. The final average concentration across all of region *i* is then the weighted average of these concentrations at *t*_*f*_ and the unchanged initial concentration *C*_*i*_(0) = *C*_*ik*_(0) from the unoccupied sites of region *i*. It is given in Eq 5.

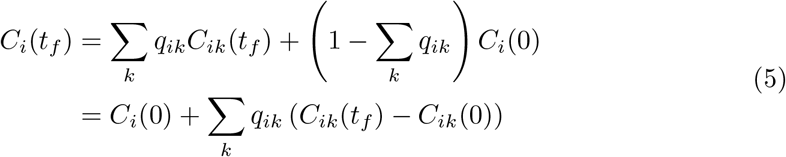

In Eq 4, it is possible that the uptake and secretion rates vary by cell type, e.g. *u* = *u*_*k*_. In our model, we assume that proliferating and quiescent cells do not differ in these parameters.

In the hybrid method, we perform the same steps as in the global method, but due to the spatial resolution we keep through diffusion, we can then apply average the extracellular concentrations for each (region, type) pairing to that specific set of lattice sites rather than averaging across each region using Eq 5.

When we consider a quorum factor, we assume that cells secrete this factor at a constant rate independent of any internalized concentration and that there is no need to track cellular uptake. Thus, we call this an export process, borrowing nomenclature from PhysiCell, and it is described simply by Eq 6. We use the explicit solution of this equation to update the concentration of the quorum factor at all lattice sites containing a cell after solving the exchange equation and before moving on to the intracellular signaling. In the global method, these contributions are averaged across each region.

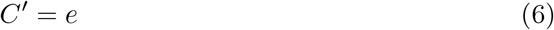

### Intracellular signaling

Intracellular signaling describes any reactions that occur with at least one molecular state variable that is unique to that cell, for example cell surface receptors or proteins in a signaling pathway (Fig 1E). Since oxygen is used by cells in aerobic respiration, that is the intracellular signaling in our model. Because our goal is to demonstrate the viability of the global method, we take a simplified approach to this well-established chemical reaction, assuming that all cells have a constant amount of glucose and so the intracellular ODE is just in the single state variable, *I*, representing the internalized oxygen concentration. It is given in Eq 7.

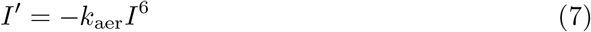

where *k*_aer_ determines the rate of this reaction and thus combines both the assumed constant glucose concentration and the actual rate of the reaction. While this equation is separable, we do not solve it with that method because that does not generalize to most intracellular reactions. We instead use the direct Euler method. While the local method solves Eq 7 once per cell, the global and hybrid methods solve this once for each (region, type) pair.

### Solving the global method in a single ODE

With the global method, it is possible to solve the molecular dynamics for a given substrate with a single ODE. The state variables are the mean internalized concentrations for each (region, type) pair as well as one concentration per region for the freely diffusing substrate. This makes for a total of *n*_regions_ × (*n*_cell types_ + 1) state variables. All of the equations describing the global method are summed up with one exception. The effects of cellular exchange are continuously averaged across the entire region rather than making temporary concentrations for the diffusing substrate accessible to each cell type. This allows for accurate PK and diffusion dynamics which are being updated at the same time as the exchange dynamics. To accomplish this, we use Eq 8. As above, *i* indicates values in region *i, k* indicates values associated with cell type *k*, and *q*_*ik*_ is the proportion of region *i* that is occupied by cell type *k*.

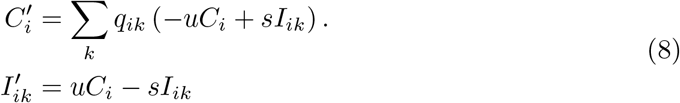

### Quiescence

After each time the molecular dynamics are solved, all cells are checked to see if their internal oxygen levels result in a phenotype change. We do this by comparing the internal oxygen concentration against a threshold value, *I*_thresh_. Any proliferating cells below this threshold are relabeled as quiescent; quiescent cells above the threshold are relabeled as proliferating.

We also consider the case where quiescence is treated as an event (like proliferation, movement, and apoptosis) rather than being determined by the internalized oxygen concentration. For this, we assume that above a threshold *I*_max_, a cell will not become quiescent, so the quiescence rate is 0. Below a threshold *I*_min_ *< I*_max_, a cell has a constant, maximal quiescence rate of *q*_max_. For *I*_min_ *< I < I*_max_, the quiescence rate is linearly interpolated between these two value so that it looks like a ramp-down function.

## Results

### Quiescence as an intracellular threshold

To show the accuracy and speed of the global method, we begin using both the local and global methods to simulate a growing group of cells on a 100 × 100 lattice over five days with 8 samples of each method. Oxygen enters the microenvironment from the bottom boundary where we assume the only relevant vasculature is. Using this information, we set up the global method so that the regions are level sets of the distance from lattice points to the blood vessel, i.e. each region is a horizontal strip one cell width wide (see Fig 1A). In solving the four differential equations in the global method, we solve them sequentially in the same manner as we do the local method. All the differential equations in the global method are solved using a matrix exponential except the intracellular signaling for which we use direct Euler. The same methods are used for the local method except the diffusion equation is solved using an LOD method.

The agreement between these two methods on cell counts is apparent throughout the simulation and they never differ in average by more than 5% in the proliferating compartment (Fig 2AC) or by more than 10% in the quiescent compartment (Fig 2BD). On internalization of oxygen by cell type, the agreement is even stronger with the relative difference being bounded by 4% and 2% for proliferating and quiescent cells, respectively (Fig S2 FigA-D). We also show that the concentration of oxygen throughout the microenvironment is similarly preserved by the global method (Fig S2 FigEF). This agreement is further corroborated by the cellular events underlying the population dynamics (Fig S3 Fig).

**Fig 2.**
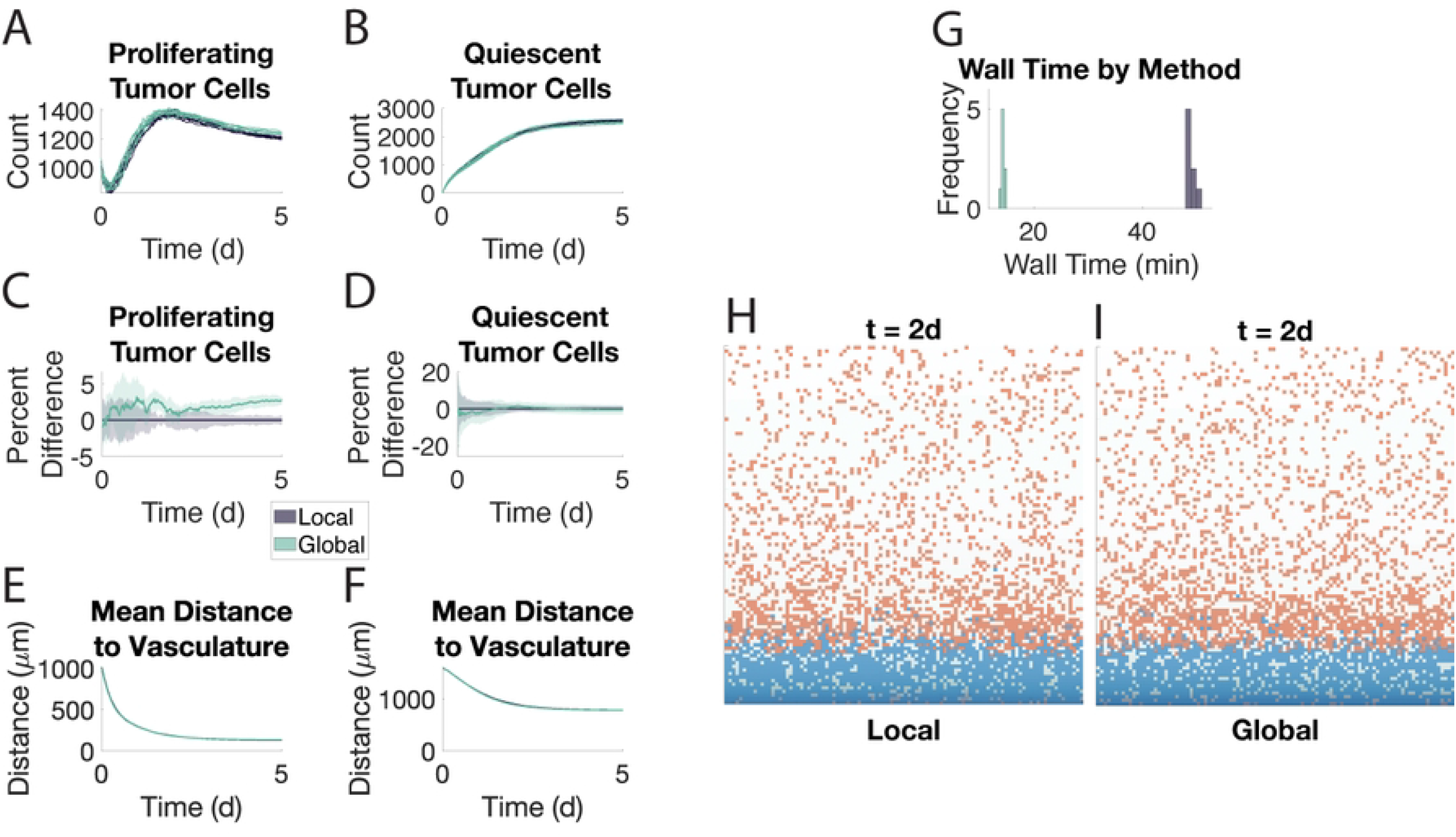
The global method agrees with the local method. Eight samples of each method are shown. Shaded area in A-F represents ±1 SD from the mean. A. Proliferating cell population in both methods. B. Quiescent cell population. C-D. Percent difference from the average population in the local method. E-F. Mean distance of proliferating (E) and quiescent (F) cells from the vasculature at the bottom of the microenvironment. G. Wall time of each method when using direct Euler to solve the intracellular signaling ODEs. H. Wall time of each method when using a Runge-Kutta method to solve the intracellular signaling ODEs for only 2 minutes. I-J. Snapshots of the local (I) and global (J) methods at *t* = 2 d. Proliferating cells are in blue, quiescent cells are in orange.

The spatial arrangements of the two cells types are in agreement between the two methods in terms of average distance to vasculature (Fig 2EF). One difference between the two methods that can be observed in comparing Figures 2IJ is that the regions in the global method are more homogeneous than the corresponding locations in the local method. That is, a single rectangular strip in the global method is much more likely to be all proliferating or all quiescent (Fig S4 Fig). When we look at the relative codensities, however, we see that this difference does not translate into a difference in proximity to the other cell type (Fig S5 FigA).

When we look at the wall times for these simulations, we see that the local method takes 5 times as long as the global method (Fig 2G). If we use MATLAB’s ode45 instead of the matrix exponential to solve all the ordinary differential equations in both methods, the local method takes 350 times as long as the global method (Fig 2H). Even if ode45 is used just for the intracellular signaling, the local method still takes 334 times longer (Fig S6 Fig).

### Quiescence as an event

To test if the global method can capture the intra-region heterogeneity observed in the local method (Fig S4 Fig) while still maintaining accuracy, we change quiescence so that it is an event cells can undergo, analogous to proliferation, death, and movement. In this way, quiescence is a stochastic process that is not solely determined by the region a cell occupies. We see that the two methods have similar levels of agreement as before, though the global method maintains a 5% difference in the proliferating compartment through Day 5 (Fig 3). The snapshots at Day 2 show noticeable cellular heterogeneity within certain regions for both methods (Fig 3HI). Indeed, the two methods show very similar intra-region heterogeneity (Fig S4 Fig) and the relative codensities of each cell type remain in agreement as well (Fig S5 FigB).

**Fig 3.**
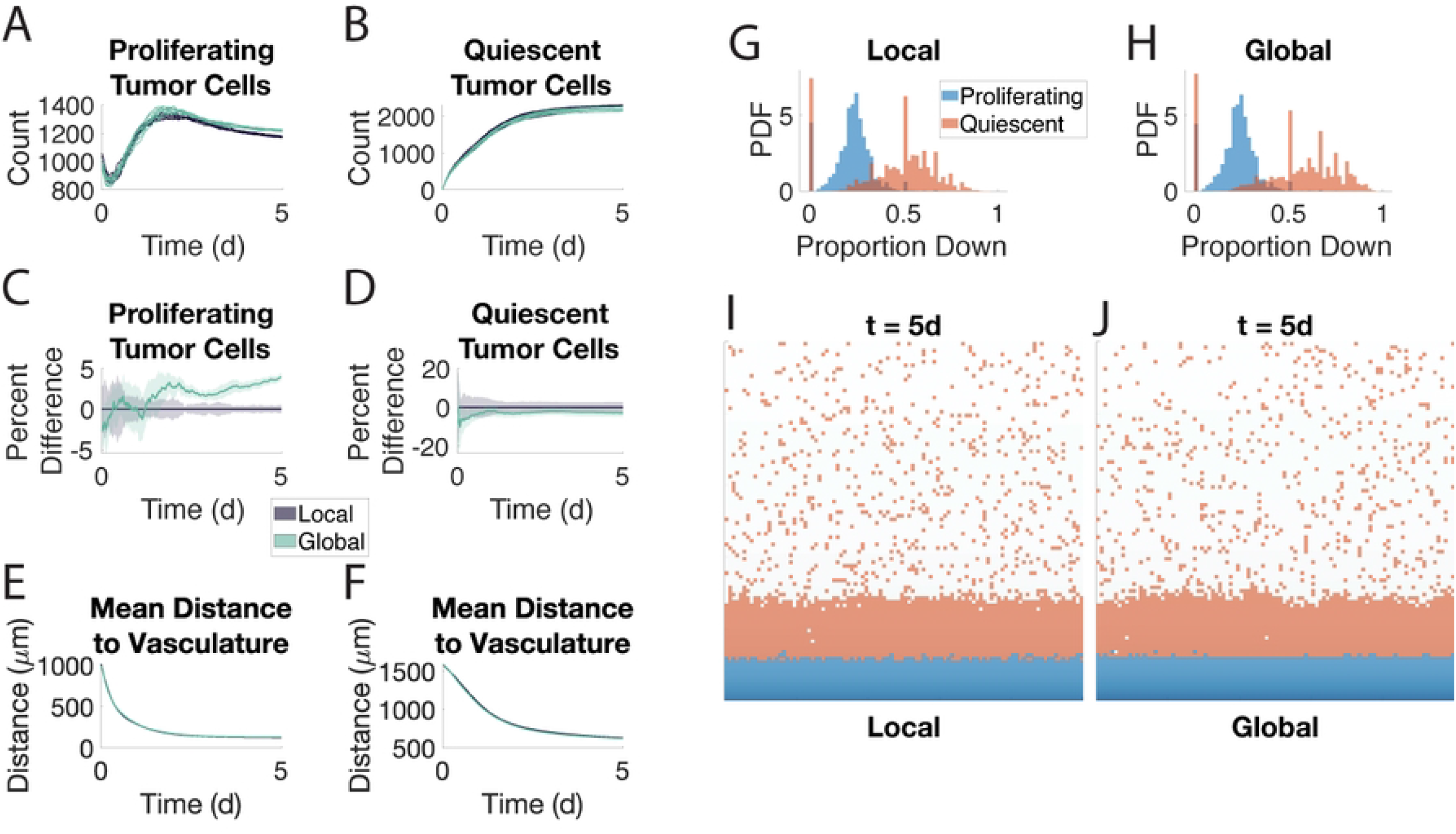
The global method agrees with the local method when quiescence is modeled as an event. Eight samples of each method are shown. Direct Euler was used to solve the intracellular signaling ODE. See caption for Fig 2.

For all of these setups, we can also use the hybrid method. While the hybrid method is slower than the global method, it still outperforms the local method (Fig S7 FigG), maintains accuracy at the macroscale (Fig S7 FigA-F), and also captures some of the microscale dynamics around individual cells (Fig S7 FigIJ).

### Chemotaxis

To test the validity of the global method in a context where cells can move along gradients at the molecular level, we allow for the quiescent cells to chemotax along the oxygen gradient. Across all our metrics, we continue to see agreement between the two methods (Fig 4A-F). This includes the quiescent compartment migrating en masse towards the blood vessel at the bottom (Fig 4F). Note how the average distance approaches 500 µm in both methods whereas previously the average distance was closer to 1000 µm (Fig 2F and Fig 3F). This is because the quiescent compartment shows similar preferences for moving along the oxygen gradient (Fig 4GH). In these simulations, the global method is three times faster than the local method (Fig S8 FigA). The intracellular signaling ODEs were solved using the direct Euler method.

**Fig 4.**
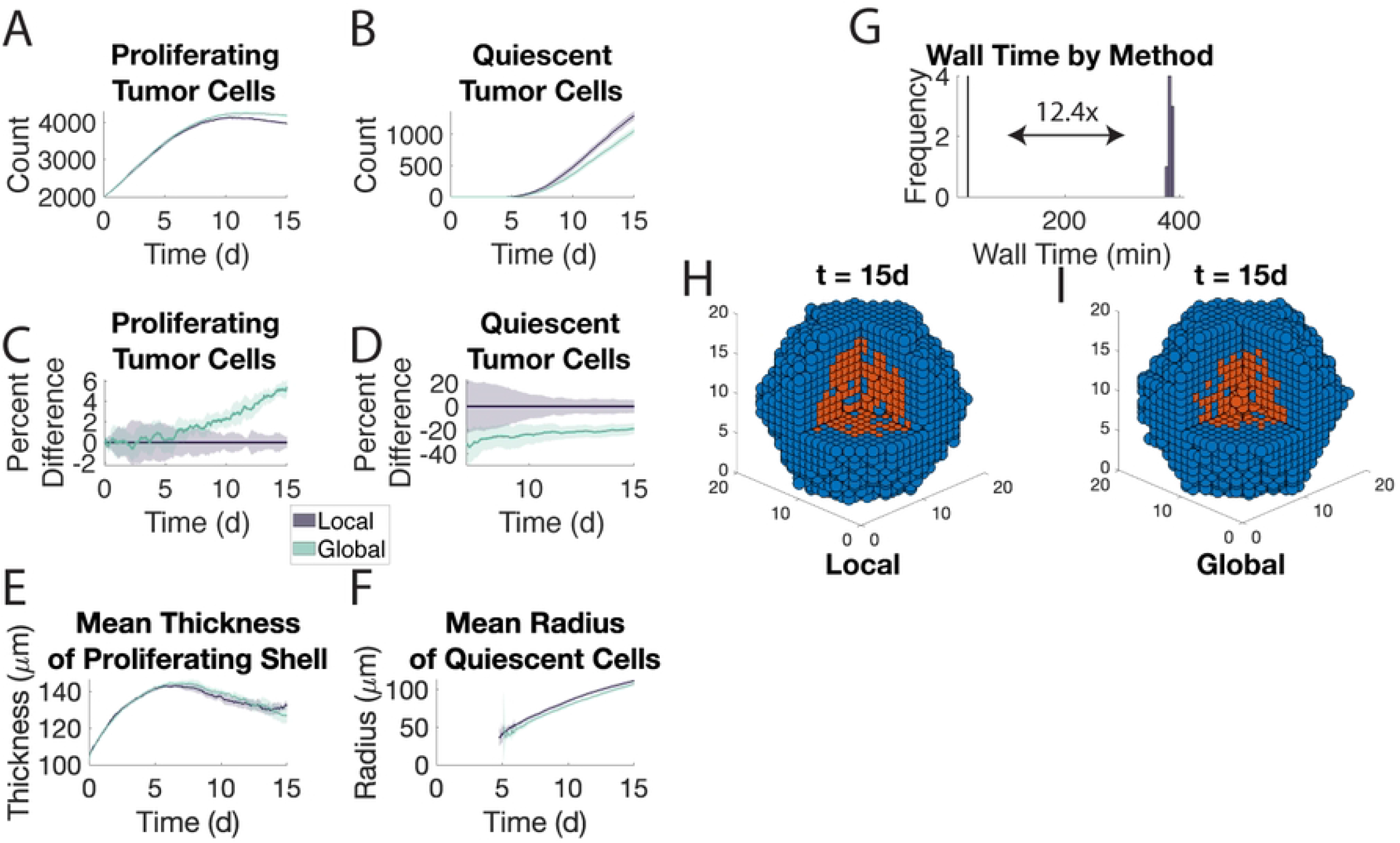
The global method agrees with the local method when quiescent cells chemotax along the oxygen gradient. Eight samples of each method are shown. Direct Euler was used to solve the intracellular signaling ODE. A-F. See caption for Fig 2. G-H. Distribution of all proportions of moves that are along the oxygen gradient by type in the local (G) and global (H) method. Specifically, for each sequence of moves an agent performs while in a single state, the proportion of those moves along the gradient is computed. These proportions are concatenated across all eight samples. Note: spikes in the quiescent histograms correspond to rational numbers with small denominators, e.g., 2*/*3 and 3*/*4. I-J. Snapshots of the local (I) and global (J) methods at *t* = 5 d. Snapshots are given at a later time point to show the effect of chemotaxis. Proliferating cells are in blue, quiescent cells are in orange.

### Quorum factor

To continue adding complexity to the model in an effort to further test the global method, we allow proliferating tumor cells to respond to a secreted substrate we call a quorum factor by chemotaxing along that gradient. This will cause the proliferating tumor cells to cluster together. The molecular dynamics for this quorum factor only include diffusion, degradation, and export by both proliferating and quiescent cells. We also implement this in a 3D microenvironment where the vasculature is assumed to be a spherical shell surrounding the center of the microenvironment. In these simulations, we initialize the agents in the center of the microenvironment to grow a spheroid. Because of these modeling assumptions, we define regions in the global method for oxygen dynamics based on the distance from the center of the microenvironment. In so doing, we partition in the microenvironment into 60 concentric spherical shells–the 3D analogue of annuli–with the outermost shells being disconnected because they would otherwise extend beyond the microenvironment box. For quorum factor, because we know from the running the local method that the agents will form a spheroid, we also use concentric shells for regions when solving the quorum factor dynamics. We only present simulations that used the global method to solve the quorum factor dynamics, choosing to continue focusing on the differences in the local and global methods in solving the oxygen dynamics. It is entirely possible to simulate the model with a using different methods for different substrates.

The two methods produce qualitatively similar results though the global method produces slightly more proliferating cells and fewer quiescent cells (Fig 5A-D). When we look at the simulations at Day 15 (Fig 5HI), we see that the spheroids look qualitatively similar despite the difference in cell counts. We also compare the radius of the quiescent core as wells as the thickness of the proliferating shell between the two methods. These are common features of tumor spheroids, a model commonly used in studies of cancer. Not only do we see a similarity from the snapshots in Fig 5HI, but we also see that the two methods agree across the simulation (Fig 5EF).

**Fig 5.**
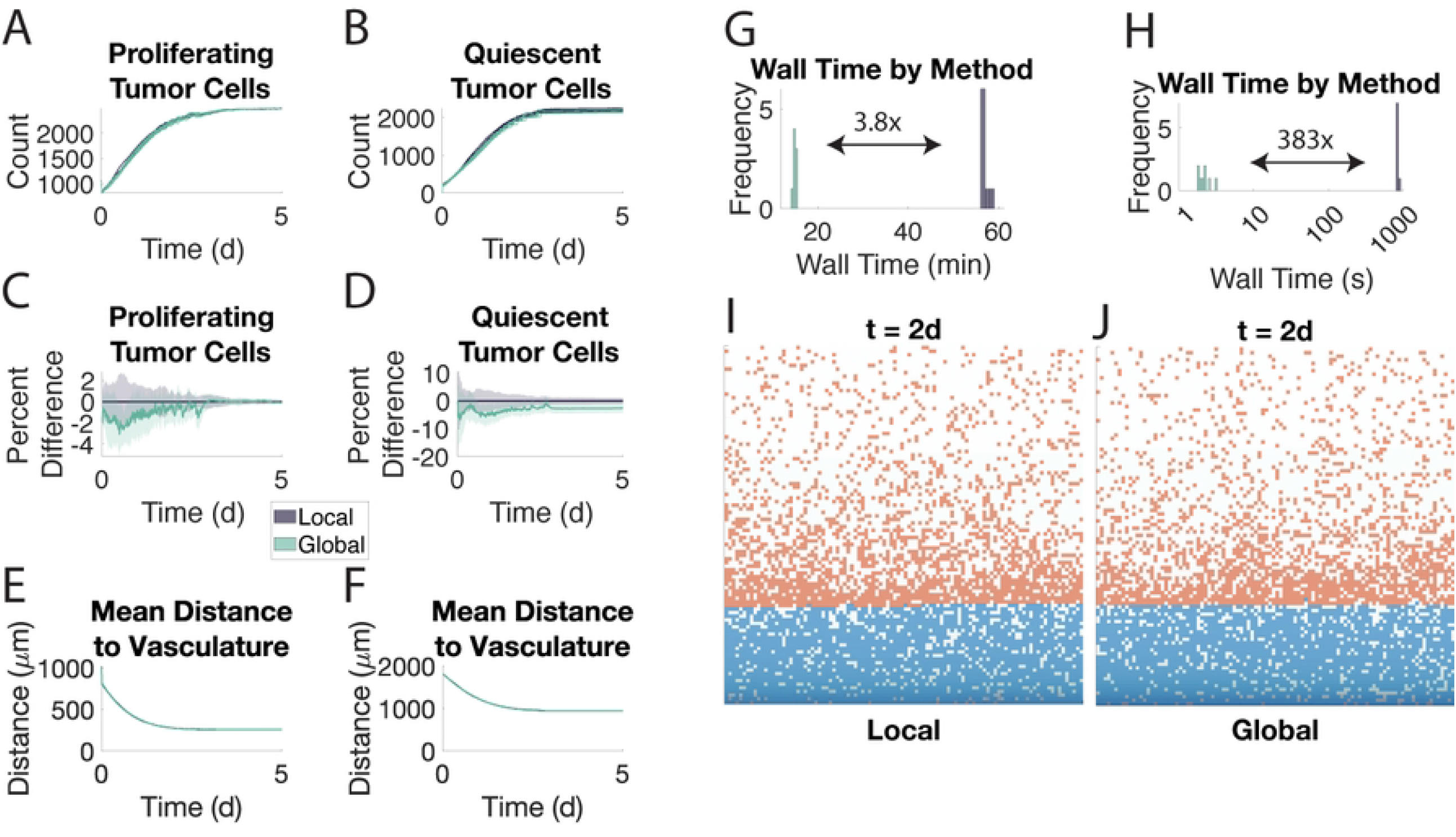
The global method agrees with the local method when vasculature surrounds the microenvironment in 3D. Eight samples of each method are shown. Direct Euler was used to solve the intracellular signaling ODE. See caption for Fig 2. E. Mean thickness of the proliferating shell. Measured by computing the difference in maximal and minimal distances of proliferating cells from the center of the microenvironment within each subset of a partition of the microenvironment and using a weighted average thereof. F. Mean distance of quiescent cells from the center of the microenvironment.

The simulations using the global method were on average 12 times faster than those using the local method (Fig 5E). Recall, that this difference is only attributable to using the global method for oxygen dynamics as the quorum factor dynamics are always solved using the global method. Furthermore, the difference in cell counts noted above means the global method has more events to simulate because quiescent cells neither proliferate nor undergo quiescence. Quiescent cells also have slower apoptosis and movement rates. Finally, the intracellular signaling dynamics were solved using direct Euler; a Runge-Kutta method or other more sophisticated ODE solver would result in an even greater difference in wall time.

## Discussion

We have shown here that the global method for simulating ABMs is sufficiently robust to handle discrete effects of the molecular scale on the cellular scale. In doing so, we extend the class of ABMs for which this method can prove useful. We also demonstrated that the global method can successfully capture the process of chemotaxis along a gradient, even when the dynamics for that substrate are solved using the global method.

The global method has previously been shown to work well in the context of continuous changes to cell fate decisions [8]. Though continuous changes need not be understood as small changes, they are nonetheless categorically different from a common occurrence in cellular biology: change of state. This adds a level of complexity to the modeled system that the global method needed to be carefully crafted to address. As cells change states, the processes they engage in and the rates at which they undertake these processes can change. Thus, a natural question for the global method is whether it can successfully capture these changes in its coarse-grained version of the system.

We took one of the most fundamental biologically processes that results in a discrete change to cell states, aerobic respiration, and used that as our paradigm for phenotype switching. This process is well-studied and of interest across fields in biology as production of energy is essential for life. By creating an ABM in which proliferating cells switch to a quiescent state in a hypoxia-dependent manner, we showed that the global method quickly and accurately reproduces the local method.

When relying on a direct Euler method for solving the nonlinear intracellular signaling ODEs, the global method outperformed the local method with approximately a fourfold speedup. The speedup magnified to over 300-times when we switched to an adaptive Runge-Kutta method, which many nonlinear ODEs would require for an accurate solution. We acknowledge that the task at hand, solving intracellular signaling ODEs for each cell, is embarrassingly parallel, opening up the possibility for parallel computing to reduce this speedup. However, the common concern for simulation time of an ABM is not around how long a single simulation requires, but how long an entire cohort of simulations requires. This is because many samples from the solution space must be drawn to understand the distribution imposed by the stochastic effects. These cohorts, in turn, are often one of many for any number of higher-level goals such as parameter estimation or therapy design. Thus, even though parallel computing can reduce the wall time of a single simulation, it still uses computing resources that could otherwise be spent solving another simulation at the same time.

By looking at both the growth dynamics and the spatial distribution of cells, we observe that the global method produces accurate simulations with respect to the local method. Across most of the metrics we look at, the percent difference between the two methods does not exceed 10%. Even in cases where it does, the qualitative behavior between the two methods is strikingly similar. We have included videos of the simulations with the online version that the reader can compare. That the two methods have qualitative agreement means that similar conclusions can be drawn using either method. For example, sensitivity to specific parameters will be preserved between the two methods, even if the they differ quantitatively.

We also explore how cell state-dependent chemotaxis affects the accuracy of the global method. Chemotaxis is a common feature of many ABMs with molecular dynamics. It also adds an additional level of interplay between the molecular and cellular levels that affects agent location. This creates a feedback loop in which the oxygen gradient affects the agent location, which then affects the oxygen gradient near the agent at its new position. We see that the global method closely follows the local method in the resulting motion of the cells.

We finally show that the global method works in the context of phenotype switching for 3D simulations with vasculature surrounding the growing spheroid. The global method reproduced the two layers of the growing spheroid from the local method: a hypoxic core and a normoxic outer shell. Thus, the global method can capture geometries that are not linear.

The global method still has room for improvements and future innovations to be adaptable to a wider range of phenomena. As one example, the parameters governing cells can vary from cell-to-cell within a given cell type as is the case in studies that include evolution [13–15]. If these parameters affect the cellular exchange or the intracellular signaling modules of the molecular dynamics, then the mean dynamics of these processes depend on the distribution of state variables as well as parameters. One possible approach would be to incorporate stochastic signaling in the global method in order to mimic the variability of cell signaling within the microenvironment. These perturbations can be derived from experiments that quantify the variability of cell signaling in a tissue both spatially and temporally. For example, if data showed that a particular signal was dependent on distance from blood vessels but with quantifiable variability within regions defined by their distance from blood vessels, then the output of the global method can be perturbed to match these observations.

Another avenue for innovation is handling two substrates that are involved in the same intracellular signaling reaction. One obvious solution presents itself: if we insist that the regions for one of the substrates is a refinement of the regions for the other, then we can solve the intracellular reactions using the finer regions. A clear corollary to this is to use the regions for one substrate to refine those of the other, which can be done for any choice of regions corresponding to the relevant geometry for any substrate. However, this can lead to large numbers of regions, slowing down the global method and reducing its utility. Worse yet, if one substrate is solved with the local method, then the intracellular signaling must be performed per cell. Can we adapt the global method to these conditions to maintain both the speed and accuracy of the global method?

One last direction for further understanding the accuracy of the global method we will discuss here is contact-mediated intercellular signaling. While we explored in this work how the global method can work as cells switch between two phenotypes, the physical interaction between neighboring cells is limited to blocking both movement and proliferation. However, cell-cell interactions are far richer than this alone. In particular, immune cell interactions with each other and with antigen-expressing cells leads to specialized behaviors that can be modulated by diffusing substrates. How does the global method perform when, for example, an immune cell and a cancer cell respond to separate diffusing substrates that both change the interaction?

## Conclusion

The global method speeds up all aspects of research involving ABMs: from model development through to model predictions. It can be used to build intuition about an ABM one is building before performing computationally expensive runs using the local method. We have demonstrated the ability of the global method in several typical use cases for ABMs. There are still more phenomena to be considered in this framework. Beyond those discussed above, cancer alone provides the examples of interstitial fluid pressure, enhanced permeability and retention, tortuous vasculature, and inhomogeneity of the extracellular matrix [16–18]. Cellular interactions with the extracellular matrix (ECM) and draining lymph nodes can be appended onto this list as well. We will continue demonstrating the utility of the global method in more and more of these cases and thus opening up this tool for more and more people to take advantage of.

## Supporting information

**S1 Fig. Oxygen concentrations in the local method suggest regions to use in the global method**. All panels show a snapshot from a simulation without any cells after a steady state has been achieved. A. Blood vessels are located on the bottom of the 2D microenvironment. B. Blood vessels are located along the entire boundary of the 2D microenvironment. C. Blood vessels are located in a spherical shell centered in the microenvironment. Concentrations are shown within this shell. The first octant relative to the center of the microenvironment has been cut away to show the concentration in the interior.

**S2 Fig. Oxygen concentrations in simulations relating to Fig 2**. Shaded areas represents ±1 SD from the mean. A. Average internalized oxygen concentration in proliferating cells. B. Average internalized oxygen concentration in quiescent cells. C-D. Percent difference between the two methods in A-B above, respectively. E. Average oxygen concentration throughout the microenvironment. F. Percent difference between the two methods in E.

**S3 Fig. Comparison of events in simulations relating to Fig 2**. A. Number of proliferations. B. Percent difference between the two methods in A. C. Number of contact inhibitions. D. Percent difference between the two methods in C. E. Number of apoptotic events

**S4 Fig. Comparison of the intra-regional heterogeneity in the local and global methods**. Each panel shows a heatmap of the composition of all regions throughout the simulation and across all samples. The *x*-axis indicates how many proliferating cells are in the region. The *y*-axis indicates how many quiescent cells are in the region. The heatmap is restricted to only regions that contained at least one of both cell types, otherwise the values along the axes would dominate the values shown here. Top row: quiescence is determined based on a threshold value for the internalized oxygen. Bottom row: quiescence is an event that cells can stochastically undergo based on internalized oxygen.

**S5 Fig. Comparison of the relative codensities in the local and global methods**. Each set of axes shows the codensity of one cell type (in the column) relative to another (in the row) for both the local and global methods. Shaded area represents ±1 SD from the mean. A. Quiescence modeled as a threshold. B. Quiescence modeled as an event.

**S6 Fig. Comparison of wall times between the two methods when solving the intracellular signaling with MATLAB’s** ode45 **and every other differential equation as described in Table 2**. Simulations were ran until *t* = 2 min. One sample was run for each method.

**S7 Fig. Comparison of the local and hybrid methods**. Compare to Fig 2.

**S8 Fig. Extended results of simulations related to Fig 4**. A. Wall time distributions for the two methods. B. Comparison of movements downward (along oxygen gradient) against all movement for both cell types across both methods. Each point represents all movements within one continuous time window in which a agent is in a constant state. C. Heat maps of the four scatter plots shown in B.

## Acknowledgements

This work was supported by NIH/NCI U01CA243075 (TLJ).

## Notes

### Competing Interest Statement

The authors have declared no competing interest.

## References

1. Dhurjati P, Mahadevan R. Systems biology: the synergistic interplay between biology and mathematics. The Canadian Journal of Chemical Engineering. 2008;86(2):127–141.

2. Ghaffarizadeh A, Heiland R, Friedman SH, Mumenthaler SM, Macklin P. PhysiCell: An open source physics-based cell simulator for 3-D multicellular systems. PLoS computational biology. 2018;14(2):e1005991.

3. Del Monte U. Does the cell number 109 still really fit one gram of tumor tissue? Cell cycle. 2009;8(3):505–506.

4. Montagud A, Ponce-de Leon M, Valencia A. Systems biology at the giga-scale: Large multiscale models of complex, heterogeneous multicellular systems. Current Opinion in Systems Biology. 2021;28:100385.

5. Gong C, Ruiz-Martinez A, Kimko H, Popel AS. A Spatial Quantitative Systems Pharmacology Platform spQSP-IO for Simulations of Tumor–Immune Interactions and Effects of Checkpoint Inhibitor Immunotherapy. Cancers. 2021;13(15):3751.

6. Fadai NT, Baker RE, Simpson MJ. Accurate and efficient discretizations for stochastic models providing near agent-based spatial resolution at low computational cost. Journal of the Royal Society Interface. 2019;16(159):20190421.

7. Lima EA, Faghihi D, Philley R, Yang J, Virostko J, Phillips CM, et al. Bayesian calibration of a stochastic, multiscale agent-based model for predicting in vitro tumor growth. PLoS Computational Biology. 2021;17(11):e1008845.

8. Bergman D, Sweis RF, Pearson AT, Nazari F, Jackson TL. A global method for fast simulations of molecular dynamics in multiscale agent-based models of biological tissues. iScience. 2022; p. 104387.

9. Widmer DS, Hoek KS, Cheng PF, Eichhoff OM, Biedermann T, Raaijmakers MI, et al. Hypoxia contributes to melanoma heterogeneity by triggering HIF1α-dependent phenotype switching. Journal of Investigative Dermatology. 2013;133(10):2436–2443.

10. Habib P, Slowik A, Zendedel A, Johann S, Dang J, Beyer C. Regulation of hypoxia-induced inflammatory responses and M1-M2 phenotype switch of primary rat microglia by sex steroids. Journal of Molecular Neuroscience. 2014;52(2):277–285.

11. Liu K, Fang C, Shen Y, Liu Z, Zhang M, Ma B, et al. Hypoxia-inducible factor 1a induces phenotype switch of human aortic vascular smooth muscle cell through PI3K/AKT/AEG-1 signaling. Oncotarget. 2017;8(20):33343.

12. Vuillefroy de Silly R, Ducimetiére L, Yacoub Maroun C, Dietrich PY, Derouazi M, Walker PR. Phenotypic switch of CD8+ T cells reactivated under hypoxia toward IL-10 secreting, poorly proliferative effector cells. European journal of immunology. 2015;45(8):2263–2275.

13. Robertson-Tessi M, Gillies RJ, Gatenby RA, Anderson AR. Impact of metabolic heterogeneity on tumor growth, invasion, and treatment outcomes. Cancer research. 2015;75(8):1567–1579.

14. Damaghi M, West J, Robertson-Tessi M, Xu L, Ferrall-Fairbanks MC, Stewart PA, et al. The harsh microenvironment in early breast cancer selects for a Warburg phenotype. Proceedings of the National Academy of Sciences. 2021;118(3):e2011342118.

15. West J, Rentzeperis F, Adam C, Bravo R, Luddy KA, Robertson-Tessi M, et al. Tumor-immune metaphenotypes orchestrate an evolutionary bottleneck that promotes metabolic transformation. bioRxiv. 2022;.

16. Jain RK, Stylianopoulos T. Delivering nanomedicine to solid tumors. Nature reviews Clinical oncology. 2010;7(11):653.

17. Shipley RJ, Chapman SJ. Multiscale modelling of fluid and drug transport in vascular tumours. Bulletin of mathematical biology. 2010;72(6):1464–1491.

18. Thurber GM, Schmidt MM, Wittrup KD. Antibody tumor penetration: transport opposed by systemic and antigen-mediated clearance. Advanced drug delivery reviews. 2008;60(12):1421–1434.

